# Shift of the insoluble content of the proteome in aging mouse brain

**DOI:** 10.1101/2022.12.13.520290

**Authors:** Cristen Molzahn, Erich Kuechler, Irina Zemlyankina, Lorenz Nieves, Tahir Ali, Grace Cole, Jing Wang, Razvan F. Albu, Mang Zhu, Neil Cashman, Sabine Gilch, Aly Karsan, Philipp F. Lange, Jörg Gsponer, Thibault Mayor

## Abstract

Aging and protein aggregation diseases are inextricably linked. During aging, cellular response to unfolded proteins are believed to decline which results in diminished protein homeostasis (proteostasis). Indeed, in model organisms, such as *C. elegans*, proteostatic decline with age has even been linked to the onset of aggregation of proteins in wild-type animals. However, this correlation has not been extensively characterized in aging mammals. To reveal the insoluble portion of the proteome, we analyzed the detergent-insoluble fraction of mouse brain tissues after high-speed centrifugation by quantitative mass spectrometry. We identified a cohort of 171 proteins enriched in the pellet fraction of older mice including the alpha crystallin small heat shock protein. We then performed a meta-analysis to compare features among distinct groups of detergent-insoluble proteins reported in the literature. Surprisingly, our analysis revealed that features associated with proteins found in the pellet fraction differ depending on the ages of the mice. In general, insoluble proteins from young models (<15 weeks) were more likely to be RNA-binding, more disordered and more likely to be found in membraneless organelles. These traits become less prominent with age within the combined dataset, as proteins with more structure enter the pellet fraction. This analysis suggests that age-related changes to proteome organization lead a specific group of proteins to enter the pellet fraction as a result of loss of proteostasis.

## Introduction

Protein homeostasis (proteostasis), crucial for the prevention of protein aggregation, has been demonstrated to decline with age and disease [1]. In order to ascertain the details of this decline, intense interest has focused on the composition of the proteins most affected by it, proteins that aggregate with age.

In *C. elegans*, loss of proteostasis was found to occur early in adulthood, coinciding with reduced heat shock response [2]. Subsequently, protein aggregates begin to accumulate [3]. In the case of the long-lived *daf2* mutant, however, more protein aggregates suggesting a protective link between aging and aggregation. Importantly, the proteins found in the insoluble fraction of the long-lived strain contained more charged residues, were more disordered and had lower aggregation scores as compared to aggregates obtained from the wild-type strain [3]. These findings suggest that there is a transition in the type of proteins that aggregate as a result of loss of proteostasis. While *C. elegans* studies have formed an important basis for aging research, they lack the important context and tissue specificity most directly relevant to aggregation-linked proteinopathies such as neurodegenerative diseases (NDs).

Chaperone, proteasome and autophagy activity have all been shown to decline in brain tissue in region and cell-type specific ways. During aging in wild type mice, levels of TRiC/CCT are reduced in neural stem and progenitor cells and in the cortex and striatum, which coincides with an increase in aggregate staining in these regions [4]. Normally, misfolded proteins are degraded by the ubiquitin-proteasome system. However, reduced proteasome activity has been measured in rodent brain tissue such as the cortex and hippocampus during aging [5]. In killifish brain tissue, reduced proteasome activity with age was shown to lead to disruption of protein complex stoichiometry and contribute to the aggregation of orphaned proteins [6]. Protein aggregates too large for proteasomal degradation are targeted by the other major route of degradation, autophagy. Detergent-insoluble ubiquitin-positive aggregates accumulate with age in *drosophila* after a decrease in autophagy-related gene expression [7]. The dysfunction of these cellular quality control components appears to contribute directly to the accumulation of aggregates with age.

There has been intense work linking biomolecular condensates (also referred to as membraneless organelles or MLOs), aging, and NDs [8, 9]. Several ND-associated mutations accelerate the vitrification of protein-rich granules [8]. Condensates are hypothesized to form through liquid-liquid phase separation creating protein-rich and dilute phases [10]. Proteins found to phase-separate are typically enriched with intrinsically disordered regions [11] allowing these proteins to form multiple, reversible, contacts creating a protein interaction network within the protein-rich phase. Age-related changes and the decline in protein quality control machinery responsible for the dissolution of condensates may increase their susceptibility to form aggregates. Formation and dynamics of stress granules has been shown to rely on ATP-dependent processes such as helicase activity and chaperonins which may decline with age [12]. Additionally, senescent cells, which accumulate with age in mice, show decreased ability to maintain stress granules [13, 14]. Together, these findings have led to the hypothesis that aggregates form within protein rich granules. Analyzing the composition of age-associated aggregates and probing changes in the properties of these protein-rich granules relative to the dilute phase of the cytosol, may reveal insights into how aggregation might begin.

Multiple proteins associated with NDs form detergent-insoluble aggregates, which can be separated by centrifugation [15]. This fractionation method has been applied extensively to identify co-aggregating and disease-associated proteins. The composition of aggregates can be used to reveal insights into the mechanisms of disease [15]. Therefore, we performed high-speed centrifugation and mass spectrometry (MS) to identify proteins that accumulate in the triton-insoluble fraction in brain tissues of old mice. We then combined this data with datasets from six other published studies using mice at various ages and disease stages. Analysis of protein features reveal a shift in the composition of proteins enriched in the pellet fraction between young and old mice.

## Results

### Identification of reduced solubility of proteins during aging

To identify proteins that aggregate with aging, we used high speed centrifugation and label-free MS based on data-independent acquisition (DIA). As proteins misfold and become incorporated into high molecular weight aggregates, they will sediment at lower centrifugation speeds. Therefore, proteins that remain in the supernatant from young mice, but enter the pellet fraction upon aging likely do so due to aggregation (Figure 1A). For this analysis, we collected cortex tissue of C57B/6 mice aged 15 and 100 weeks, and analyzed the composition of supernatant and pellet fractions. Using DIA-MS we quantified 4,385 protein groups. As expected, these protein groups clustered strongly by fraction, and to a lesser extent by age (Figure S1A). Samples showed strong correlation within fraction groups (Pearson correlation > 0.96 for supernatant and > 0.89 for pellet) and less strong correlation across groups (Pearson correlation 0.82-0.88) (Figure S1B). Proteins that become insoluble with aging were identified by comparing the composition of the pellet fractions. A group of proteins (171) was found to be enriched in the pellet fraction of the older mice (t-test p-value < 0.05 and log_2_(fold change) > 0.5) (Figure 1B, C). GO analysis reveals this group of proteins is enriched for terms known to be affected by aging such as mitochondria and lysosomes (Figure 1D) [16, 17]. A smaller group of 59 proteins was identified as more insoluble in the pellet fraction from young mice (t-test p-value < 0.05 and log_2_(fold change > −0.5) (Figure 1B). This group of proteins was enriched for terms such as histones, PDZ domains, lipid degradation and ND pathway-associated proteins (Figure 1D). Analysis of the supernatant fraction revealed minimal changes in soluble protein abundance between the two age groups (Figure S1C), consistent with previous findings [18]. Importantly, proteins that are enriched in the pellet fraction with age tend to also be more abundant (Figure S1D). However, for a majority of these proteins (148/171) this enrichment is not statistically significant. Given that proteins are frequently expressed at their limit of solubility, we chose to proceed with the analysis of all 171 proteins with the hypothesis that an increase in abundance, even minor, would cause the proteins to aggregate [19, 20].

**Figure 1.**
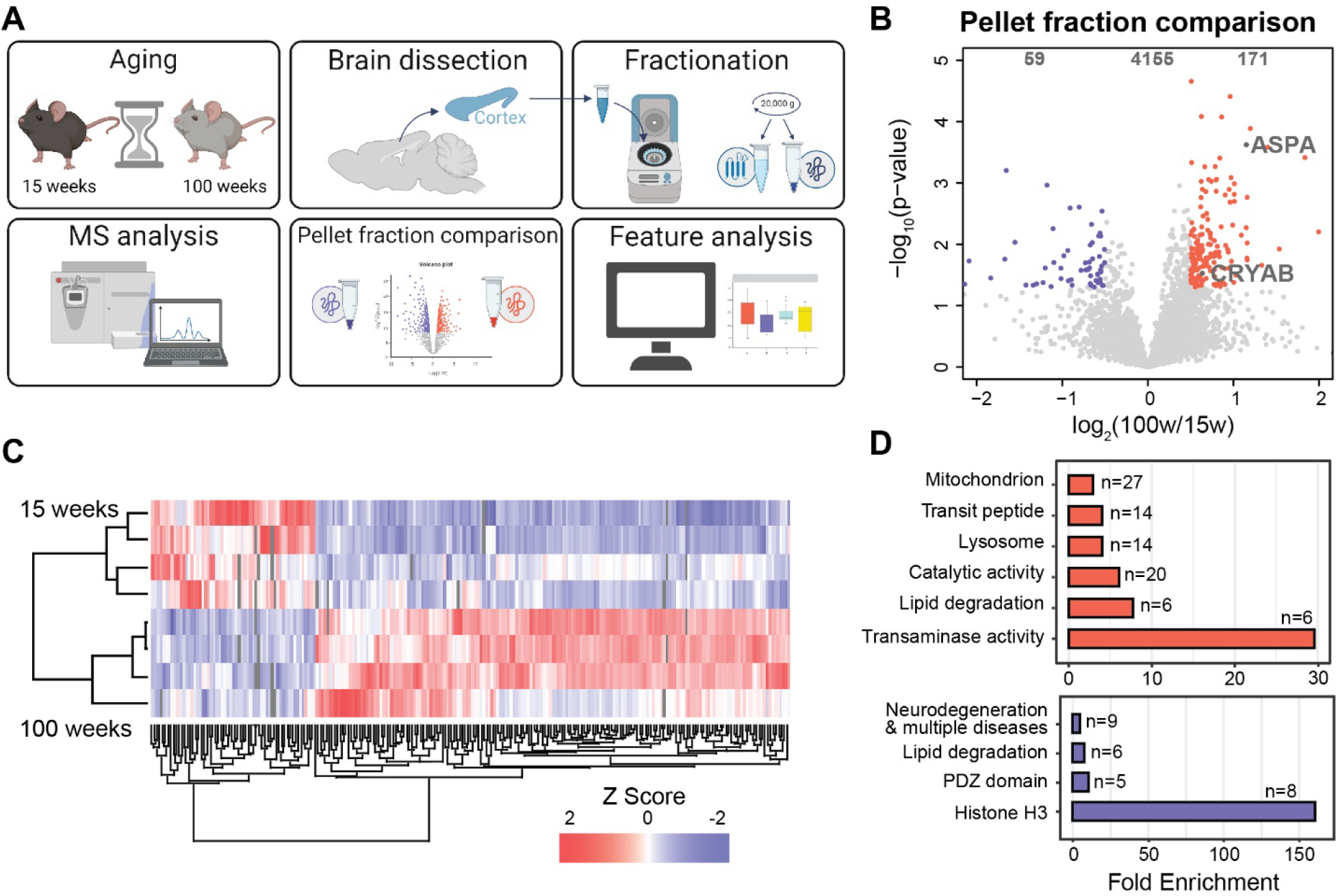
Detergent insoluble proteins accumulate in the brain tissue of 100-week-old mice. A) Outline of the experimental workflow. B) Comparison of the composition of detergent insoluble fractions obtained from the cortex of 15 and 100-week-old (n=4, n=4) mice using a two-sample paired t-test. C) Euclidean distance clustering of z-scored intensities from the significantly altered proteins. D) Top GO terms (Benjamini-Hochberg corrected p-value < 0.05) for proteins enriched in the pellet fraction of old mice (pink) and young mice (purple). Numbers of protein groups in each category are indicated. Created with BioRender.com

To confirm the aggregation of these proteins, cellulose acetate filter trap assays (FTAs) were performed. Two candidate proteins were selected for validation based on their enrichment in the pellet fraction including: the small heat shock protein alpha crystallin B chain (CRYAB) and aspartoacylase (ASPA), which has previously been implicated in the early onset ND Canavan disease (Figure 1C) [21]. In the FTAs, both candidate proteins show significantly increased immunodetection in the three old mice as compared to the three young mice (Figure 2A, B). Old mice show a greater increase in signal for ASPA than for CRYAB, consistent with their enrichment in the mass spectrometry data. In the total tissue lysate, levels of ASPA also increase. However, this increase is less than the increase in signal observed by FTA and is not significant (Figure 2C). CRYAB shows no change in abundance in the total tissue lysate (Figure 2D). This observation suggests that the increase in signal observed in the FTA is not due to changes in expression level. Importantly, while both proteins were enriched in the triton insoluble fraction and FTA in old mice, we could not see evidence for increased deposition of these proteins into foci by microscopy (Figure S2B-C), indicating our fractionation method may enable the early detection of events linked to the age-associated proteostasis collapse.

**Figure 2.**
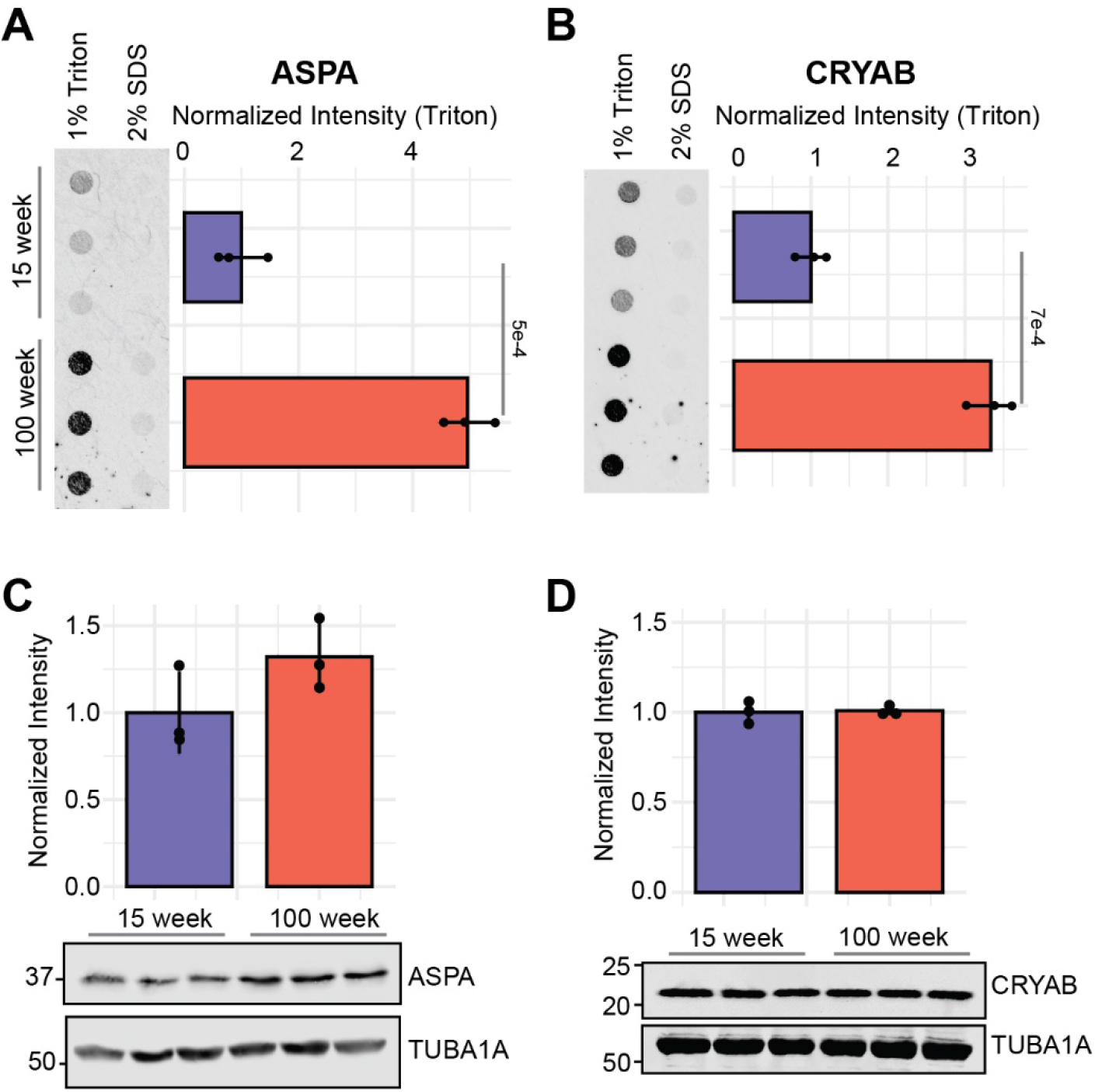
Validation of the aggregation of candidate proteins. FTA of A) ASPA and B) CRYAB using the cortex of 15 and 100-week-old mice (n=3, n=3). The FTA signals are normalized to the mean of the corresponding young data points. Western blot signal from C) ASPA and D) CRYAB is normalized the □ - tubulin loading control. Intensity ratios are then normalized to the mean of the young data points. Standard deviations and p-values of two-sample paired t-tests are indicated.

### Meta-analysis of aging and ND models

Centrifugation-MS is a widely used technique that has been applied to several ND models and cases. We therefore sought to compare our analysis to other studies that have used different ND mouse models [22–27]. Selected studies used a variety of brain regions, extraction and fractionation methods, and MS approaches. For each dataset, we required on the order of 100 proteins enriched in the pellet fraction in a given condition using the enrichment cutoffs outlined by the publication. The majority of the proteins are identified in only one or two datasets, whereas less than 11% of the proteins were identified in three or more independent analyses (Figure S3A). A comparison of the enriched proteins from each mouse dataset grouped by age or disease showed that there is very little overlap in the proteins that pellet in each condition (Figure S3B-F). This finding suggests that there is not a specific group of proteins that are universally affected by aging or disease. With very few shared proteins between disease models or ages, we proceeded with a protein feature analysis to determine whether these proteins, may, instead share a set of characteristics that would make them susceptible to aggregation.

To identify features associated with aging and NDs, we collected 38 protein features, similar to properties that we previously assessed when characterizing proteins affected by stress response [28–30]. Using principal component analysis (PCA), relevant features were identified by removing features until >80% of the variance was captured in dimension 1. It was found that the remaining protein features separate the datasets by age. Datasets obtained from mice less than four months (11-13 weeks) are well separated from mice over 18 months (87-104 weeks), which include our new dataset (Figure 3A). The other datasets (26-52 weeks) are less well separated, although they more often cluster with datasets derived from older mice. Together, this finding suggests that, despite a high variability in protein composition, the age of the mice affects the composition of the pellet fraction, whose proteins share a set of features; regardless of the disease model, tissue type, or extraction method. The features that most strongly contribute to the variance in the first dimension of the PCA are associated with protein structure, solubility, and RNA-binding (Figure 3B).

**Figure 3.**
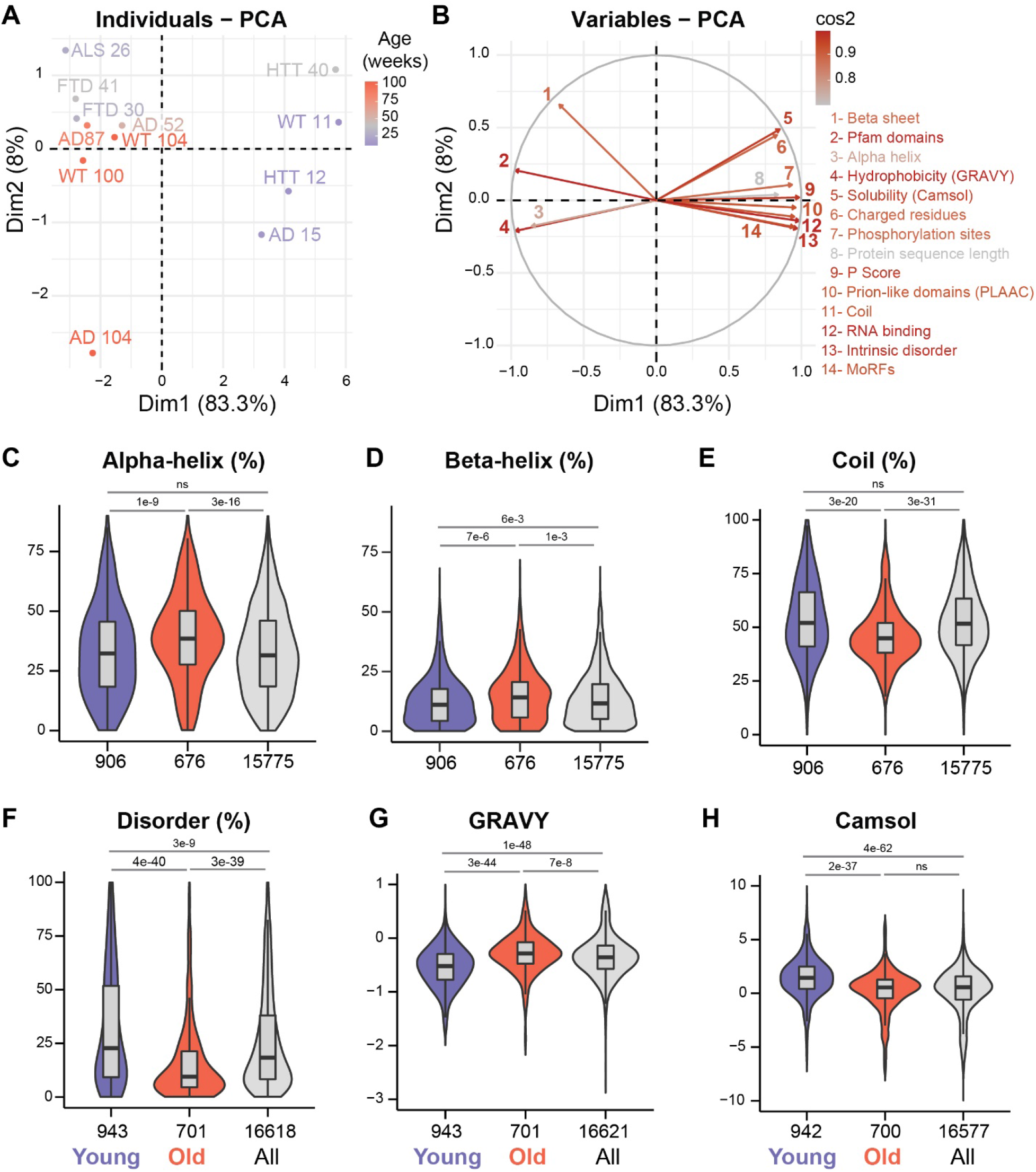
Protein feature analysis reveals features that separate datasets by age. A) PCA analysis of the 12 different data set. B) Features that were used in the PCA and their contribution to the distance in the PCA. C-H) Violin plots comparing the distribution of the datasets binned into young (<15 weeks; purple) and old (>80 weeks; pink), and of the mouse proteome (grey) for the indicated features. P-values (Hochberg adjusted Wilcoxon test) and number of proteins assessed for a given analysis are shown.

To better understand how aging may affect the composition of the proteins in the pellet fractions, we pooled the different datasets in two groups, proteins derived from mice younger than four months and older than 18 months. Comparison of gene ontology terms present in these datasets shows enrichment for terms related to translation, RNA-binding and MLOs such as the nucleolus and nuclear speckles in the young group (Figure S4A). In the old group, mitochondrial proteins are enriched as well as proteins in pathways related to NDs (Figure S4B). Proteins in the older dataset show enrichment for secondary structure (alpha helix and beta sheet), as well as a depletion for disorder and coil content relative to both the proteome and the young dataset (Figure 3C-F). We observed similar results when comparing the proteins enriched in pellet fraction due to age in our mouse study (Figure S5A-D). In agreement, the proteins in the combined old dataset, as well as in our own mouse analysis, have more of their sequence in pfam domains (Figure S6A, S5E). However, it should also be noted that proteins in the old dataset have shorter sequences, while those in the young dataset tend to have longer sequences in comparison to the proteome (Figure S6B). By contrast, proteins in the young dataset show enrichment for intrinsic disorder and coil content while showing depletion in beta sheet content in comparison to the proteome (Figure 3D-F). Together, this result indicates that proteins in the old dataset are more structured while proteins in the young dataset contain less secondary structure and are likely intrinsically disordered.

Given the difference in structure content between datasets, we also expect differences in features associated with solubility. Indeed, proteins in the old dataset are more hydrophobic (GRAVY; Figure 3G and S5F). In contrast, proteins are less hydrophobic and more soluble (CamSol) in the young dataset Figure 3G-H). In addition, these proteins have more charged amino acids likely contributing to their lower GRAVY score (Figure S6C). Proteins in the young dataset also have more reported phosphorylation sites, which could further modulate their charge (Figure S6D). While both datasets show elevated supersaturation scores relative to the proteome, proteins from the old dataset have higher supersaturation scores compared to those from the young dataset (Figure S6E). Proteins with high supersaturation scores have previously been shown to be co-aggregated with disease-associated proteins and have been found to be enriched in the detergent-insoluble obtained from mouse brains with deficient neuronal autophagy [31, 32]. Additionally, proteins in the old dataset have a higher percentage of their sequence in aggregation-prone regions (TANGO) while those in the young dataset show reduced TANGO scores (Figure S6F)[33]. This difference likely contributes to the difference observed in supersaturation scores. Interestingly, both groups show a trend toward longer half-lives in the cell using data obtained from neurons and brain tissue (Figure S6G-H)[34, 35]. Notably, proteins in the old dataset have longer half-lives over those in the young dataset. Overall, proteins in the young dataset show enrichment for features associated with solubility while those in the old dataset show enrichment for features associated with aggregation. This is especially interesting given that they also show longer lifetimes and may be less likely tightly regulated by degradation.

### Membraneless compartment enrichment

Many of the features enriched in the young age group are also associated with proteins found in MLOs. Proteins in the young dataset are more likely to participate in pi-pi interactions (PScore) a feature that is correlated with the ability to liquid-liquid phase separate (Figure 4A) [36]. Pi stacking interactions facilitate protein-RNA interactions, so it is fitting that the young dataset is enriched for RNA-binding proteins (Figure 4B). RNA-binding proteins are thought to be important for the formation of phase-separated RNA-containing granules [37]. The young dataset also shows enrichment for prion-like domains (PLAAC) which can promote phase separation (Figure S6I) [38, 39]. We recently established several computational tools to predict localization of proteins in MLOs, such as the Membraneless organelle and Granule Z-Score (MaGS) [28, 29]. Accordingly, proteins in the young dataset have significantly higher MaGS values relative to both the proteome and the old dataset indicating that they are more likely to be found in MLOs (Figure 4C). Therefore, we next assessed whether protein components of specific MLOs were enriched in the young dataset. Using entries from the data repository of liquid-liquid phase separating proteins (drLLPS), we found enrichment for stress granules, neuronal granules, chromatid bodies, the post-synaptic density and nucleolus among the proteins in the young dataset (Figure 4D)[40]. Also enriched were proteins that have been shown to phase separate *in vitro* (i.e., categorized as droplet). In contrast, while some proteins are components of MLOs, the old dataset shows no significant enrichment for these phase-separated compartments with the exception of the post-synaptic density. Our analysis suggests that proteins which localize in granules are potentially depleted from the insoluble fraction with age, which would be counterintuitive, given the hypothesized role of phase-separated granules in the progression of aggregate formation [8, 9].

**Figure 4.**
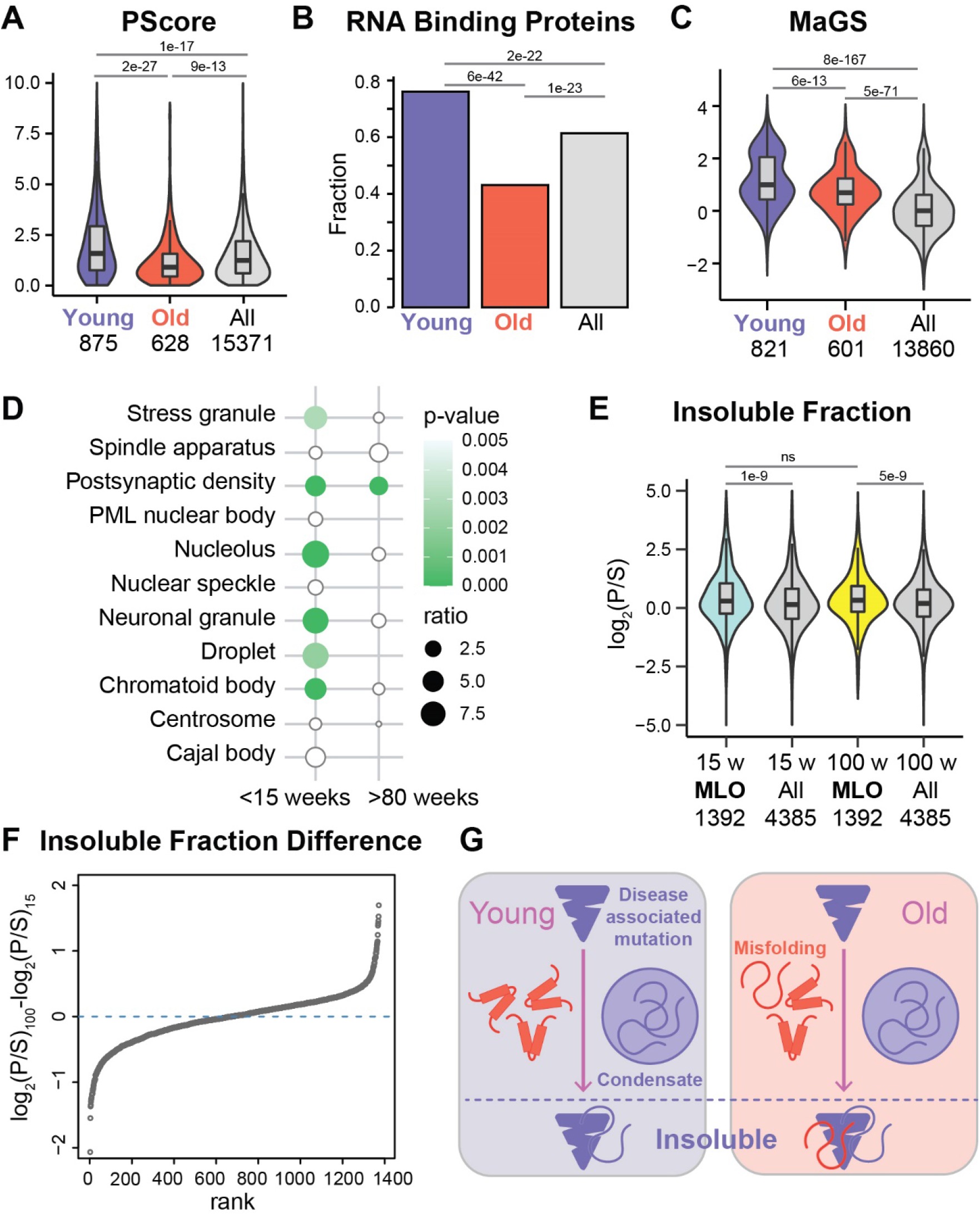
Young datasets are enriched for features and proteins associated with MLOs. A and C) Violin plots comparing A) the distribution of P-Scores and C) Mammalian granule z-scores for each protein. P-values for all violin plots were calculated using a Benjamini-Hochberg adjusted Wilcoxon test. B) Bar plot comparing the proportion of proteins predicted to be RNA-binding with Benjamini-Hochberg corrected p-values obtained from fisher test. D) Representation of Fisher test results indicating enrichment of MLO proteins as annotated by the drLLPS. Dot size represents the odds ratio and green intensity represents the Bonferroni adjusted p-values. E) Violin plots comparing the average intensity of proteins identified in the pellet and supernatant fractions (P/S) in old and young mice in our study. P-values were calculated using a Benjamini-Hochberg adjusted Wilcoxon test. Number of proteins assessed for a given analysis are shown. P/S ratio of all quantified proteins (grey), of MLO proteins in the young mice (blue) and of MLO proteins in the old mice (yellow) are shown. F) Ranked plot of the difference in pellet/supernatant ratio between old and young mice for all annotated MLO proteins. G) Schematic representation of the composition of the pellet fraction in young and old mice.

To better investigate the fate of MLO proteins, we directly examined the pellet/supernatant ratios (P/S) in our mouse study. This analysis allows us to assess which portion of a given protein is enriched in the pellet fraction [26]. Consistent with our previous results, proteins annotated to be in MLOs are significantly more enriched in the pellet fraction in young mice in comparison to all identified proteins (Figure 4E). However, protein annotated as belonging in MLOs are also enriched in the pellet fraction in old mice in comparison to all identified proteins. There was no significant difference between the P/S ratios of MLO proteins between young and old mice. Similar results are obtained when comparing the change in P/S ratio between the age groups, as we find that most MLO proteins display minor differences (log_2_(P/S)_100_-log_2_(P/S)_15_) < 1) (Figure 4F–S7A). The lack of widespread change in P/S ratios for MLO proteins suggests that proteins associated with MLOs are not depleted from the pellet fraction during aging. Therefore, the main change to the composition of the pellet fraction is the addition of other proteins that are not prominently localized in MLOs and that become more enriched in the pellet fraction as a result of age-related loss of proteostasis.

As several MLO proteins have been associated with ND, we sought to more carefully examine their fate to verify that these proteins do not become more insoluble. Notably, homologs of several MLO proteins that are associated with ND such as FUS, TDP43, HTT and other well characterized stress granule proteins (G3BP1, EIFs1, PABP1) are quantified in our proteomic analysis. The P/S ratio of these proteins indicates these proteins do not become more insoluble in the older mice (Figure S7B-D). We also performed an FTA on another well-known stress granule marker, TIA1, which was not quantified in our analysis. This marker shows signal in both young and old mice and no significant difference in signal between age groups (Figure S7E-F). This analysis shows that well characterized condensate proteins are not further enriched in the insoluble fraction upon aging.

### Aggregating proteins in human

We sought to extend our analysis to human studies in order to determine whether the trends seen in old mice translate to humans. We collected three centrifugation-MS datasets derived patient tissues including frontotemporal dementia (FTD), Alzheimer’s disease (AD) and chronic traumatic encephalopathy (CTE) cases [41–43], and also one dataset using laser-capture microdissection MS in AD patient tissues [44]. Similar to the mouse data, there is little overlap among the proteins identified in each study (Figure S8A). Nonetheless, the proteins in these datasets share similar features to those observed in the old mouse datasets, excepting the CTE dataset. Of the four datasets, three showed depletion for intrinsic disorder (Figure S8B) and an enrichment for secondary structure content, either beta-sheet or alpha helix (Figure S8C-D). All three of these datasets also showed depletion for coil regions (Figure S8E). The CTE dataset is markedly different and stands out for having no significant differences in secondary structure. While no significant changes were observed for most of the features associated with solubility, all datasets show high supersaturation scores (Figure S8F). Furthermore, three out of four datasets show significantly lower PScores than controls, suggesting the proteins in these datasets are less likely to phase-separate (Figure S8G). Finally, two of the three datasets show depletion for RNA-binding proteins (Figure S8H). The CTE dataset, notably the youngest dataset (mean age of 60), shows no significant changes in any of these features. Overall, this meta-analysis of human data confirms that more structured proteins tend to accumulate in the insoluble fraction with ND.

## Discussion

It has been widely hypothesized that the chronic stress conditions of aging and disease effect the liquid-like properties of MLOs. For example, in mouse models of tauopathy TIA1 foci accumulate with the progression disease pathology eventually colocalizing with tau foci [45]. Such research formed the basis of our initial hypothesis that we would observe accumulation of proteins with features associated with MLOs when comparing the composition of the insoluble portion of the proteome between young and old mice. Instead, we observed depletion of these features in our dataset.

During aging, a small group of proteins becomes insoluble in 1% triton in the wild-type mouse cortex. Given the solubility of both ASPA and CRYAB in 2% SDS, demonstrated by our FTA, these proteins are most likely not solid amyloid-like aggregates. This group of proteins is enriched for GO terms such as mitochondria and lysosome, both of which have been shown to decline with aging [16, 17]. Interestingly, transit peptides, sequences that target proteins to organelles, were among the enriched terms. This finding offers a potential explanation for the incorporation of mitochondrial proteins into aggregates as subcellular localization may become disrupted during aging or ND [46]. Additional enriched terms are related to metabolism such as catalytic activity, lipid degradation and transaminase activity. Comparison of this initial dataset with additional published data revealed insights into the effects of aging and disease state on aggregation.

Despite little overlap in identified proteins between ages or disease models, common protein features were discovered that are shared between protein aggregates obtained from mice of similar ages. As observed in *C. elegans* data, aggregates that accumulated in long lived worms are enriched for more charged proteins that contain more intrinsically disordered regions. This result suggests that there is a protective role for the aggregation of these proteins. Proteins enriched in the pellet fraction of the young mouse models share similar properties suggesting a role of these proteins in modulating the solubility of the proteome in stress or in the case of a disease-associate protein [3]. The elevation in MaGS and PScore values as well as the enrichment in MLO proteins suggest that the protein in this dataset readily become integrated into protein-rich compartments. These compartments may facilitate the protective sequestration of disease-associated proteins [47]. However, it is also hypothesized that incorporation into protein-rich compartments may favor the formation of aggregates over time [48]. Misfolded proteins may become incorporated into ribonucleoprotein (RNP) granules through interactions with the prion-like domain of granule proteins. Chaperone activity is required to prevent the conversion into protein aggregates [49]. During aging, the activity declines and stress response becomes less effective. It is therefore tempting to hypothesize that aggregates form through an increase in misfolded protein-nucleated RNP granules. However, our analysis suggests that there is not an accumulation of RNP granules with age in the assessed conditions.

Proteins that aggregate with age show distinct features from those in the younger models. Features associated with the formation of MLOs, such as RNA-binding, PScore, and disorder, tend to be depleted from this group of proteins (Figure 3F, 4A-B). However, there are no widespread shifts in the P/S ratio obtained from the mouse cortex for RNA-binding or MLO proteins (Figure 4E-F). Together, these findings suggest that there is not an accumulation or depletion of MLO proteins during aging in our analysis, but rather a new group of proteins enters into the pellet fraction. Recent work in *drosophila* found that RNP granules increase in size during aging, but do not display changes in Fluorescence Recovery After Photobleaching (FRAP) signal. Additionally, these granules were not observable by centrifugation suggesting they are distinct from protein aggregates [50]. While the proteins that aggregate in younger mice seem characteristic for proteins involved in MLOs, our analysis suggests that additional proteins begin to aggregate during aging due to insufficient protein folding or degradation capacity.

The enrichment for features associated with secondary structure in proteins that aggregate with age suggest that these proteins may be more dependent on chaperone activity to properly fold. Folding and refolding of proteins typically relies on ATP-dependent chaperones [51]. In the aging brain, there is a shift in expression from ATP-dependent to ATP-independent chaperones [52]. The enrichment for secondary structure in our findings, combined with the higher GRAVY score, suggest that the proteins enriched during aging may have a hydrophobic core that could make them more prone to aggregate in the event of damage or improper folding. The high supersaturation score of these proteins suggests that they may be on the edge of their solubility limit and susceptible to aggregation upon age-related perturbations in cellular homeostasis. Interestingly, we recently showed that yeast proteins with similar properties (e.g., higher beta sheet content) tend to remain thermo-sensitive for a longer period of time following their synthesis [30]. Therefore, an alternative explanation is that the accumulation of these proteins in the pellet fraction occur due to a lower folding capacity affecting proteins being translated. In both cases, the features of these proteins suggest that they are more likely to be affected by age-related shifts in chaperone levels due to their precarious solubility and likely dependence on a declining population of chaperones.

Analysis of human datasets shows greater similarity to the trends observed in the older mouse models suggesting that a similar class of precarious proteins is affected due to aging. Human datasets show depletion for disorder and generally show enrichment for beta sheet and alpha-helix structures. However, the enrichment for secondary structure content varies between the datasets. Similar to the aged mouse dataset, human datasets do not appear to show enrichment for features associated with RNP granules. Depletion in PScore was observed for three of the four datasets and depletion for RNA-binding was observed for two. Both of these features trended lower than the proteome in all of the datasets.

The diminished protein folding capacity results in a more general decline of structured proteins resulting in their aggregation due to damage or improper folding. The notable lack of increased of granule-associated properties in insoluble proteins from aged mice is surprising given the hypothesis that aggregates form through incorporation of disease-associated protein into MLOs. One possibility is that this method captures an immobile fraction of MLO proteins in the triton-insoluble pellet but, importantly, the amount of MLO proteins in the immobile fraction does not appear increase upon aging. Based on this assumption, we see no evidence of age-dependent accumulation of proteins in condensates in our experimental set up.

## Methods

### Collection of mouse tissues

This animal study was performed under UBC Animal Care Protocol #A18-0067. Two cohorts of C57BL/6J mice were analyzed; one for the MS experiment and a second for the orthogonal validations. For the MS analysis, 4 old (100 weeks), 4 adult (78 weeks) and 4 young aged-match control (15 weeks) male mice were bred and maintained in the UBC Animal Resource Center and fed a standard diet (Teklad X2920, Envigo, USA). For the orthogonal analysis (immunoblots and IF), 15 and 100-week-old male C57BL/6J mice (6 each) were purchased from the Jackson Laboratory and kept at the UBC facility for another 10 weeks. The authors acknowledge that they unfortunately did not consider the impact of mouse sex at the time of the study design. Mice were anesthetized using isoflurane gas, and perfused with 20mL of 1xPBS containing 1x HALT protease and phosphatase inhibitors through the left ventricle. For the MS and FTA experiments, cortex sections were collected and snap frozen in liquid nitrogen for storage. For IF, left brain hemispheres were fixed in 4% PFA overnight followed by cryoprotection in 20% sucrose and embedding in OCT media.

### Enrichment for triton X-100 insoluble aggregates

Detergent insoluble aggregates were extracted from the cortex of mice aged 15 and 100 weeks as described previously [53]. Tissues were dissected, snap frozen and stored in liquid nitrogen until lysis. Proteins were extracted by cryogrinding in liquid nitrogen followed by lysis in a 1% triton X-100 containing buffer with sonication. Cell debris was cleared by centrifugation for 10 min at 3,000 g at 4°C. The supernatant was moved to a new tube and the aggregated fraction was collected by centrifugation at 20,000 g for 30 minutes at 4°C. Protein pellets were washed 2x with 1mL of lysis buffer and centrifuged at 20,000 g for 10 minutes at 4°C. Pellet proteins were resuspended in 5% SDS by sonication. Concentrations of the pellet and supernatant fractions were determined by Bradford assay.

### Trypsin digest

Trypsin digestions were performed on S-trap columns (Protifi). For the pellet fraction, all of the material was loaded onto filters (about 50 μg). Aliquots of supernatant were combined 1:1 with a 10% SDS containing S-trap loading buffer and loaded onto filters (about 50 μg). Binding, washing and digestions were carried out according to the manufacturer S-trap protocol with an additional methanol chloroform wash to remove excess lipids. Trypsin was added at a 1:25 ratio and the reaction was incubated overnight at 37°C. The resulting peptides were then acidified and desalted using C18 stage tips.

### MS data acquisition and data analysis

Proteomic analysis was performed on a Q Exactive HF Orbitrap mass spectrometer coupled to an Easy-nLC 1200 liquid chromatography system (Thermo Scientific) configured with a 20nl sample loop, 30 μm ID steel emitter. Mobile phase A was 2% Acetonitrile (ACN) in 0.1% Formic acid (FA) in water and Mobile phase B was 95% ACN in 0.1% FA in water. A 50 cm μPAC^TM^ analytical column (Pharmafluidics) with trap column was used. The analytical column temperature was set to 50°C.

Spectral libraries were from pooled supernatant and pellet peptides which were fractionated by high pH RPLC into 96 fractions and concatenated into 8 fractions. After concatenation, each fraction was analyzed by LC-MS/MS via data dependent acquisition (DDA) by injecting 1 μg per sample. Samples were eluted for 85 minutes using a 60-minute chromatic gradient with a 300 nL/min flow rate (0 min: 4 %B; 5 min: 9% B; 10 min: 10 % B; 15 min: 12 %B; 20 min: 14 %B; 25 min: 15 %B; 30 min: 17 %B; 35 min: 18 %B; 40 min: 19 %B; 45 min: 21 %B; 50 min: 24 %B; 55 min: 27 %B; 60 min: 80 %B; 85 min: 80%). Pre-column equilibration was at 3 μL at a flow rate of 3 μL per min. A full MS1 spectrum (400–1,800 m/z) was collected at 60,000 resolution. Maximum injection time was at 75 ms and an AGC target value of 3e^6^ was set. The top 12 precursors were selected for fragmentation. MS2 scans were acquired at 15,000 resolution. Maximum injection time of 50 ms and AGC target value of 5e^4^ was set. Normalized collision energy was set to 28. Dynamic exclusion duration of 20.0 s was used.

For data independent acquisition (DIA) of individual supernatant and pellet peptide samples, 800 ng per sample was injected and eluted for 85 minutes using a 60-minute chromatic gradient as above. A full MS spectrum scan (300-1650 m/z) was collected with a resolution of 120,000. Maximum injection time was 60 ms and AGC target value was 3e^6^. DIA segment spectra was acquired with 24-variable window format with a resolution of 30,000. AGC target value was 3e^6^. Maximum injection time was set to ‘auto’. The stepped collision energy was set to 25.5, 27.0, 30.0.

Spectral library preparation was performed using Biognosys. Spectral library preparation was comprised of individual DIA samples and fractionated DDA samples. Mouse fasta and iRT fasta files were used as sequence data bases. Digestion type and enzyme was set to Specific and Trypsin/P, respectively, with 2 missed cleavages allowed. Minimum and maximum peptide length was set to 7 and 52, respectively. Acetyl (Protein N-term) and Oxidation (M) were set as variable modifications; carbamidomethyl (C) was set as fixed modification.

DIA data was extracted using Spectronaut and the spectral library described above using the default BGS Factory settings. The resulting protein-level intensity values were used for further enrichment analysis. While our original MS analysis included samples from 78-week old adult mice, this data was not further analyzed. Two-sample, unpaired t-tests were performed using the base R function. Equal variance was assumed for the two age groups in each comparison.

### Gene ontology analysis

Gene ontology enrichment analysis was performed using the Database for Annotation, Visualization and Integrated Discovery (DAVID 2022) using the default annotation categories. Similar terms were clustered by function and representative terms from each cluster with a fold enrichment greater than 2 and Bejamini-Hochberg corrected p-value less than 0.05, were plotted according to fold enrichment [54]. For the comparison of GO terms between datasets the clustering function was not used. Molecular function, biological process and cellular component terms were compared between groups and filtered for presence in 3 datasets from the old or young group. The resulting GO terms were manually clustered by grouping terms from the same hierarchy and selecting representative terms.

### Immunofluorescence in Tissue Sections

Frozen sagittal sections of the left hemisphere were generated by cryosectioning at the size of 12 μm using a CM 3050C cryostat (Leica, Germany) and brain tissue sections were thaw-mounted on commercially available ProbeOn Plus charged slides (Fisher, USA). Prior to staining, gelatin coated slides containing brain tissues were dried overnight at room temperature. After drying, the slides were washed twice for 5 minutes each in 0.01 M PBS. Following washing, antigen retrieval was performed by boiling in 10 mM Sodium citrate pH 6 with 0.05% Tween-20 in a pressure cooker for 30 minutes. Blocking and permeabilization was done using 5% goat serum and 0.1% triton in 1x PBS for 1 hour at room temperature in humidified chamber. Primary antibodies rabbit anti-ASPA (ab223269; 1:50) and rabbit anti-CRYAB (ab13497; 1:50) were incubated overnight at 4°C. After primary antibody incubation, the sections were washed twice for 5 minutes each and secondary antibodies, goat anti-rabbit Alexa 568 (Invitrogen A11011; 1:200), were incubated for 90 minutes at 4°C, following slides were washed twice for 5 minutes each. DAPI staining was performed for 10 minutes followed by mounting with Dako fluorescent mounting media (Molecular Probe, Eugene, OR). Images were acquired with an Olympus FV1000 microscope using a 60x objective (UPLSAPO60XW).

### Immunoblots

Mouse cortex tissue was lysed as described above. Protein concentrations were determined using Bradford assay. Cell debris was removed by centrifugation at 3,000 RCF for 10 minutes. For filter trap assays, lysates were diluted to 0.4μg/μL with either lysis buffer or with lysis buffer with 2% SDS (final concentration). A 0.2μm cellulose acetate membrane and Whatman filter paper were fitted into a Scie-Plas dotblot apparatus. Tissue lysates were loaded onto the dotblot apparatus 50μL at a time for a total of 100μL. Vacuum was applied until all liquid flowed through. Each well was washed 3x with the corresponding buffer. The membrane was then blocked in 5% milk in Tris-buffered saline with 0.1% tween (TBS-T) for 1 hour at room temperature followed by immunodetection. For western blots, 40 μg of protein from each tissue lysate was loaded on a 10% acrylamide SDS-PAGE gel. Semi-dry transfer was performed using a Bio-Rad Trans-Blot Turbo Transfer System. Membranes were blocked for 1 hour in 5% milk in TBS-T followed by overnight incubation at 4°C with primary antibodies. Secondary antibodies were diluted 1:10,000 and incubated for 1 hour at RT. The following antibodies were used: goat anti-TIA1 (SAB2501039; 1:1,000), rabbit anti-ASPA (ab223269; 1:2,000), rabbit anti-CRYAB (ab13497; 1:2,000), mouse anti-TUBA1A 1:2,000 (ab126165), donkey anti-goat (LIC-926-32214), goat anti-mouse (LiCor 926-32210, 1:10,000) and goat anti-rabbit (LiCor 925-32211, 1,10,000). Imaging was performed and quantified with the Odyssey system (LiCor).

### Feature analysis

Protein half-lives were obtained from previously published studies conducted in mice (days) and in neurons (hours) [34, 35]. PCA was performed using the FactoMineR package in R version 4.1.2. For each dataset, the mean value for each feature was calculated. PCA was then performed using the mean values for each feature in each dataset. Beginning with 38 features, the feature with the lowest contribution to the variance was removed until 80% of the variance was captured in dimension 1. The final PCA contains 16 features.

Features analyzed included: intrinsic protein disorder as calculated by DISOPRED 3, prion-like domains calculations from PLAAC, the pi-pi interaction PScore, granule localization probability, aggregation-prone region prediction in TANGO, the grand average hydrophobicity score, RNA-binding likelihood calculated by RBPPRED, protein secondary structure using SCRATCH, molecular recognition features from the MoRF ChiBi System, and protein solubility scores calculated with Camsol [28, 33, 36, 39, 55–61]. Other protein features, such as the percent composition of amino acids or protein sequence length, were calculated using in-house Perl (v5.30.0) scripts. Furthermore, database information on protein features were gathered to supplement calculations, including: integrated whole-organism protein abundance from PaxDB, observed domain melting temperatures, protein-protein interaction counts from BIOGRID, Pfam domain annotations (version 33.0 30357350), and annotated phosphorylation site information from PhosphoSite Plus [62–67]. Data processing was handled using an in-house Perl (v5.30.0) pipeline and then was plotted using automatically generated R scripts (v4.1.2) from a Bash (v5.0.17) wrapper. Data processing scripts available at (https://github.com/cmolzahn/aging_proteomics.git) and feature plots in all datasets can be found at (https://cmolzahn.shinyapps.io/meta_study_app/).

### Membraneless organelles enrichment analysis

MLO annotations were downloaded from the data repository of Liquid-Liquid Phase Separation version 1.0 April 11, 2022 (drLLPS) [40]. The data was filtered to contain entries for *Mus musculus* that included scaffold and client proteins. The entries were compared against old and young datasets and the number of proteins in each compartment was counted. This was compared against the total number of proteins in each compartment compared to a proteome size of 17,102 using a fisher test (R version 4.1.2 Stats package) follow by Bonferroni p-value correction.

### Computation evaluation of the solubility of MLO or ND-associated proteins

ND-associated proteins were obtained from NDAtlas [68]. To obtain lists of disease-associated proteins that are also annotated in MLOs, the list of annotated ND proteins was compared against proteins annotated as scaffolds in drLLPS. The mouse homologues of the resulting 10 proteins were compared against the mouse cortex data to obtain a P/S ratio.

## Acknowledgments

We thank members of the Mayor lab for discussions, the UBC bioimaging facility and the proteomics facility for help and support. This work was supported by a grant from the Canadian Institutes of Health Research (CIHR; PJT 148489 to T.M. and PJT-162131 to A.K). C.M. is the recipient of a UBC Scholarship. A.K. is Tier 1 Canada Research Chair in Myeloid Cancers.

## Figures

**Figure S1.**
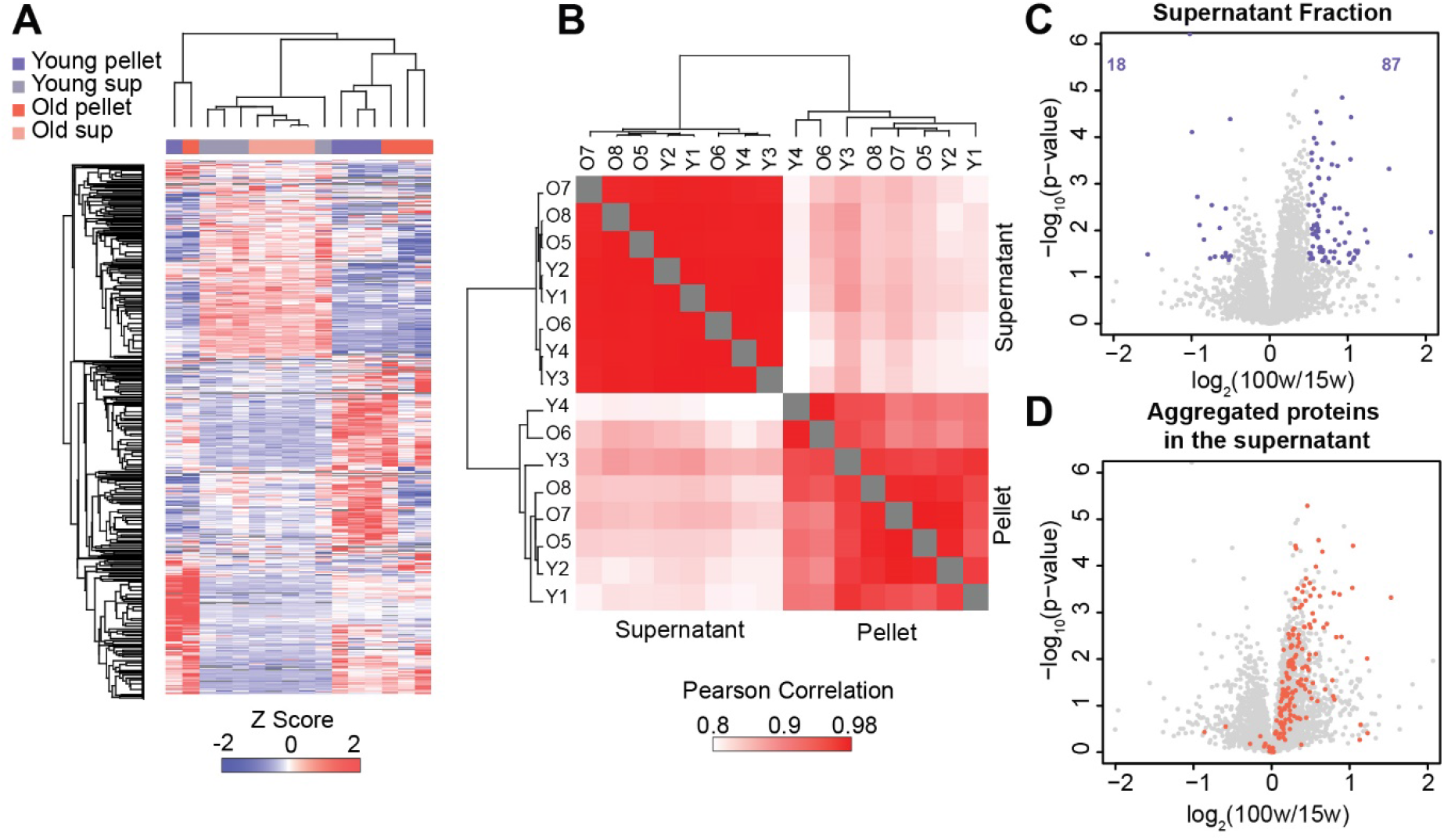
Protein level changes in abundance with age are not widespread. A) Euclidean distance clustering of z-scored protein intensities in the different indicated fractions. B) Euclidean distance clustering of Pearson correlations between each indicated fraction. C) Comparison of the composition of the supernatant fractions obtained from the cortex of 15 and 100-week-old (n=4, n=4) mice using a two-sample paired t-test identifies 105 proteins that change in abundance with age. D) Proteins that are enriched in the pellet fraction and quantified in the supernatant fraction are indicated in red in the same volcano plot shown in Figure 1B.

**Figure S2.**
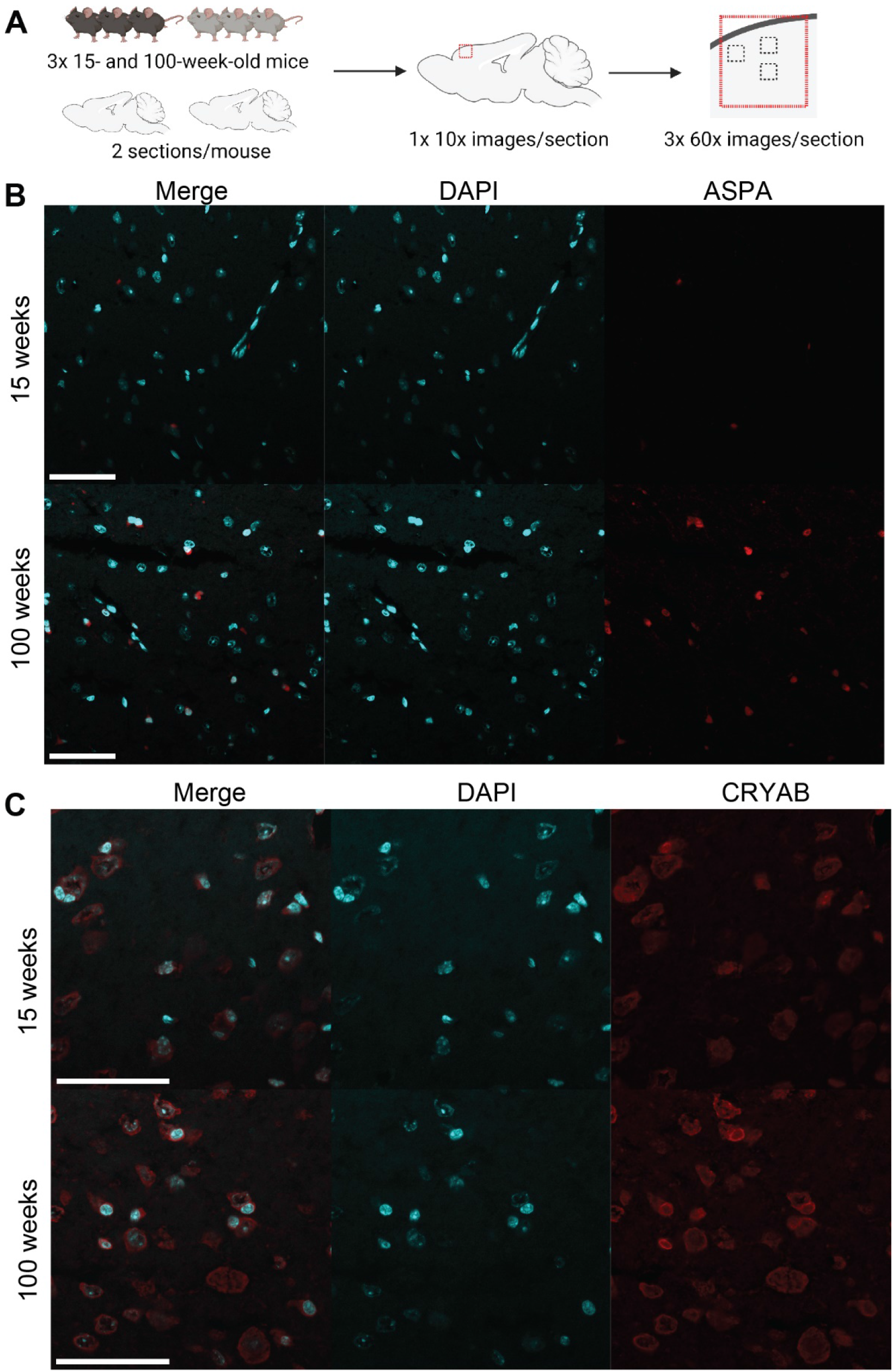
ASPA and CRYAB foci are not observed by microscopy. A) Schematic of cortex image collection in two sections obtained from 3 mice from each age. B) Representative images of B) ASPA and C) CRYAB signals in the cortex of young and old mice. While the ASPA signal appears in more cells in the old mice, it does form distinct foci. CRYAB appears diffuse in the cytosol and nucleus of young and old mice. Scale bars = 50 μm.

**Figure S3.**
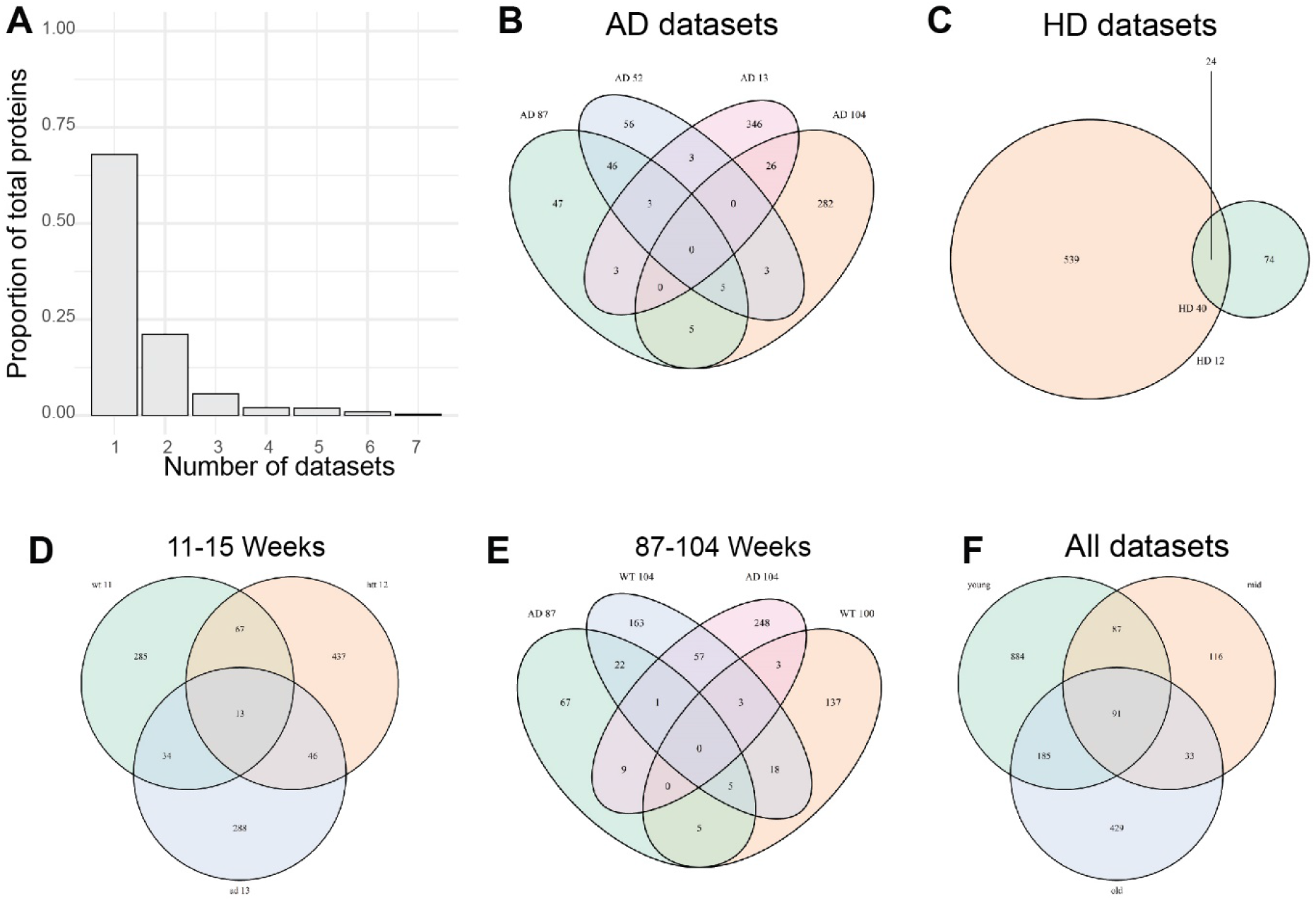
Protein solubility datasets show very little overlap. A) The proportion of proteins that are observed in the indicated number of datasets. Venn diagrams representing the overlap in B) the Alzheimer’s datasets C) Huntingtin’s datasets D) mice ≤15 weeks E) mice ≥87 weeks and F) datasets binned into young, middle and old age.

**Figure S4.**
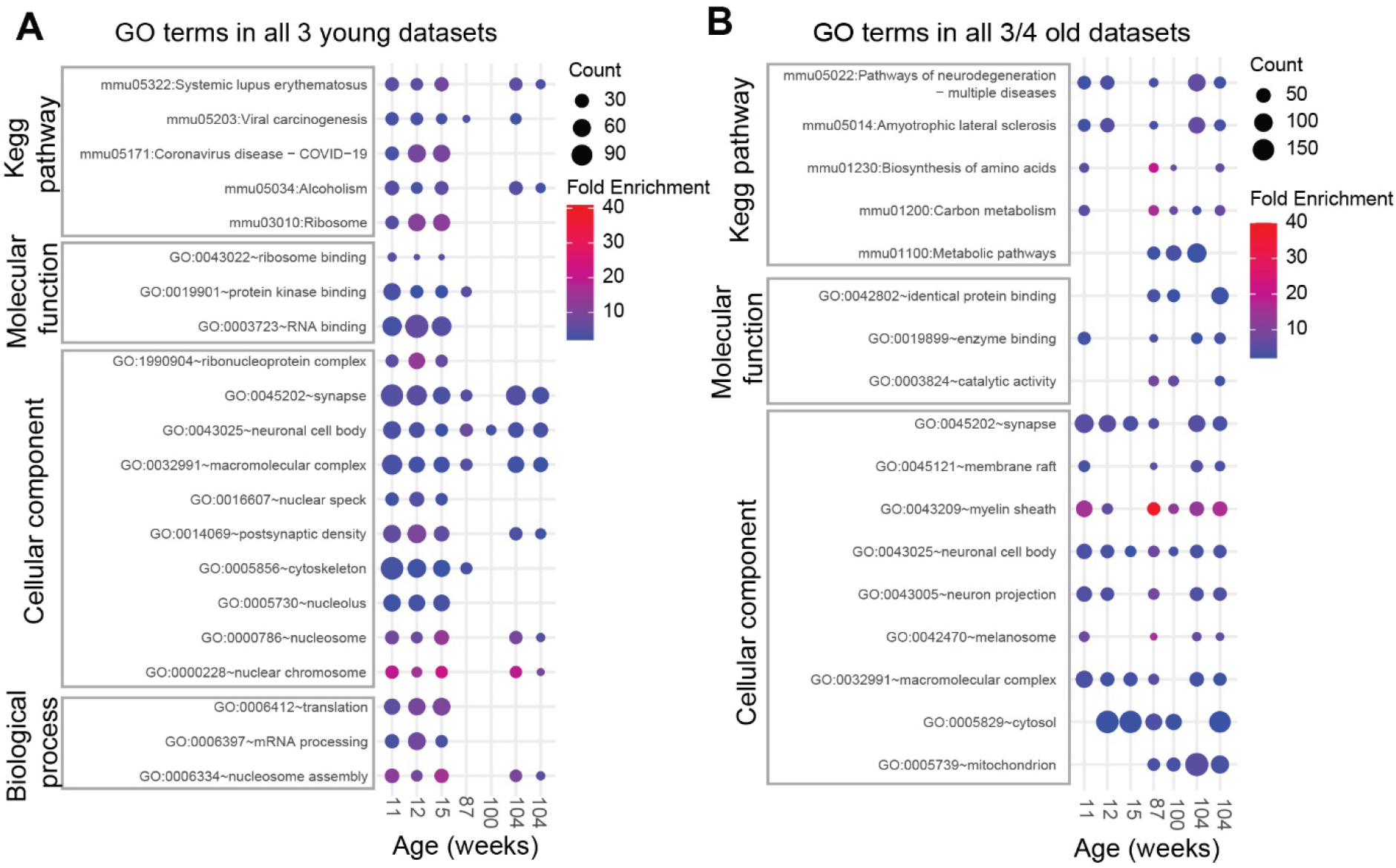
GO analysis of old and young datasets. Significantly enriched GO terms found in A) all three of the young datasets and B) three of the four old datasets are plotted by fold enrichment and the number of proteins in each group (count). GO terms were filtered for significance (FDR corrected p-value < 0.05) and then compared across datasets.

**Figure S5.**
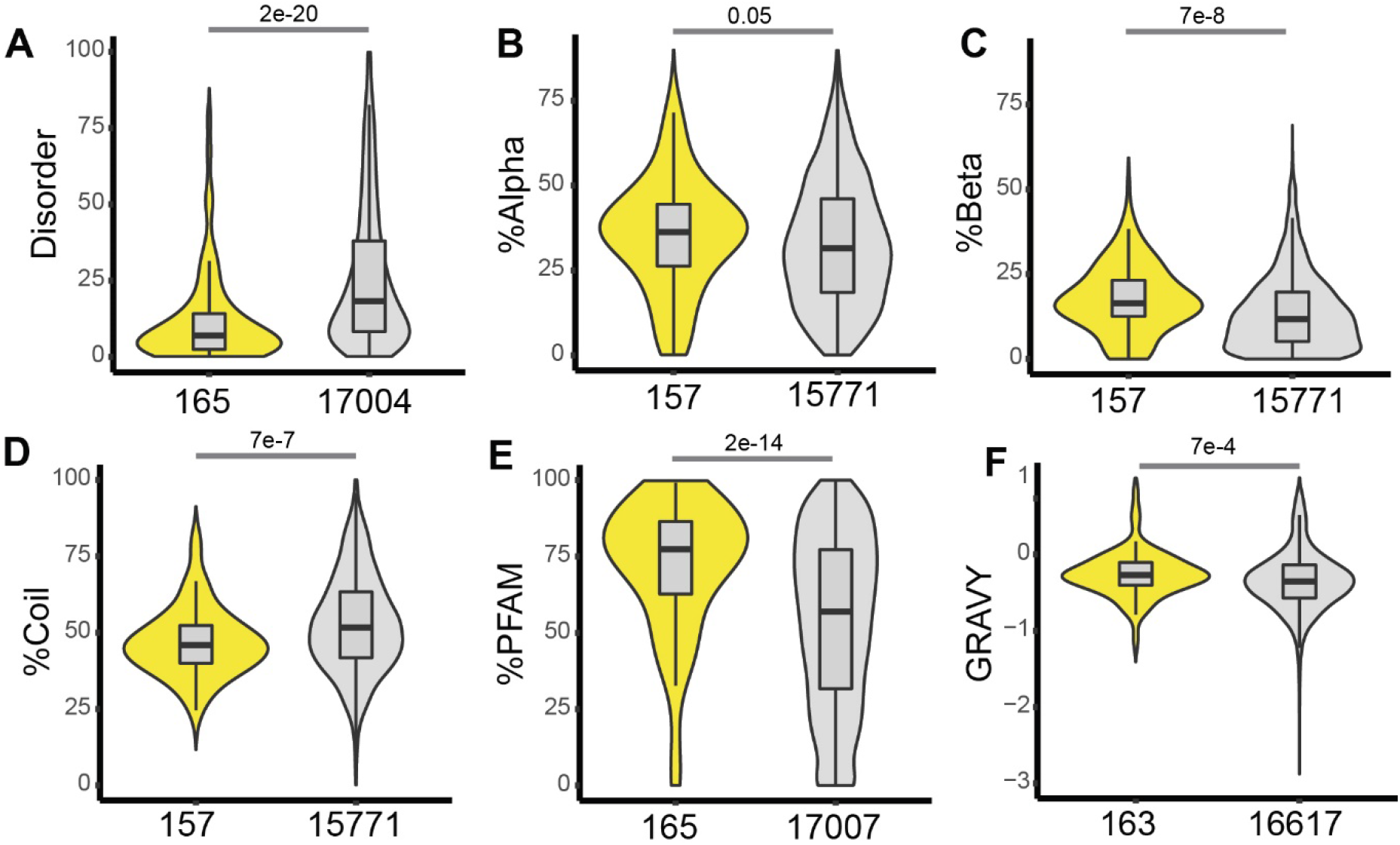
Proteins in the pellet fraction from the cortex of 100-week-old mice analyzed in this study are enriched for features associated with secondary structure. A-F) Violin plots comparing the distribution of the indicated features. A) percent intrinsic disorder; B) percent alpha helix; C) percent beta sheet; D) percent coil; E) percent of the protein sequence found in pfam domains; F) hydrophobicity score (GRAVY). Benjamini-Hochberg adjusted p-values obtained from a Wilcoxon test. Number of proteins assessed for a given analysis are shown

**Figure S6.**
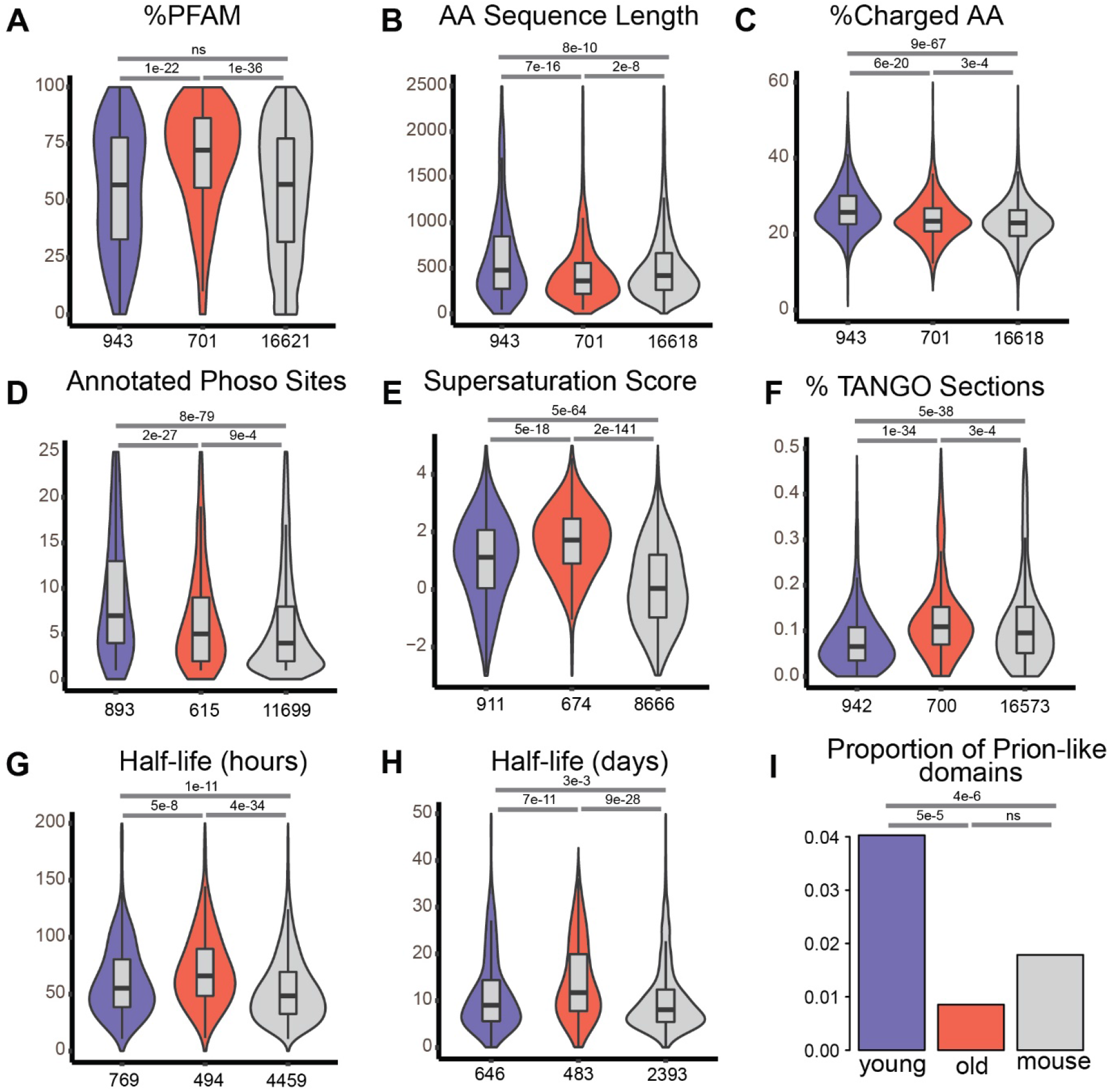
Feature analysis reveals difference in protein structure, solubility and turnover between old and young groups. A-H) Violin plots comparing the distribution of the indicated features. The following analyses are show: A) The proportion of the protein sequence in Pfam domains; B) The amino acid sequence length; C) The percentage of charged residues in each amino acid sequence; D) The number of phosphorylation sites; E) The protein’s supersaturation scores, which is a measure of abundance and aggregation propensity; F) The percentage of each amino acid sequence found in aggregation-prone regions (Tango score); Protein half-lives measured in G) hours obtained from primary neuron cell culture and H) days obtained from mouse brain tissue. Proteins in the young dataset are shown in purple, in the old dataset in pink and from the mouse proteome in light grey. The number of proteins assessed for a given analysis are shown below each bin. P-values for all violin plots were calculated using a Hochberg adjusted Wilcoxon test. I) The proportion of proteins with prion-like domains (PLAAC). FDR corrected p-values obtained from fisher test.

**Figure S7.**
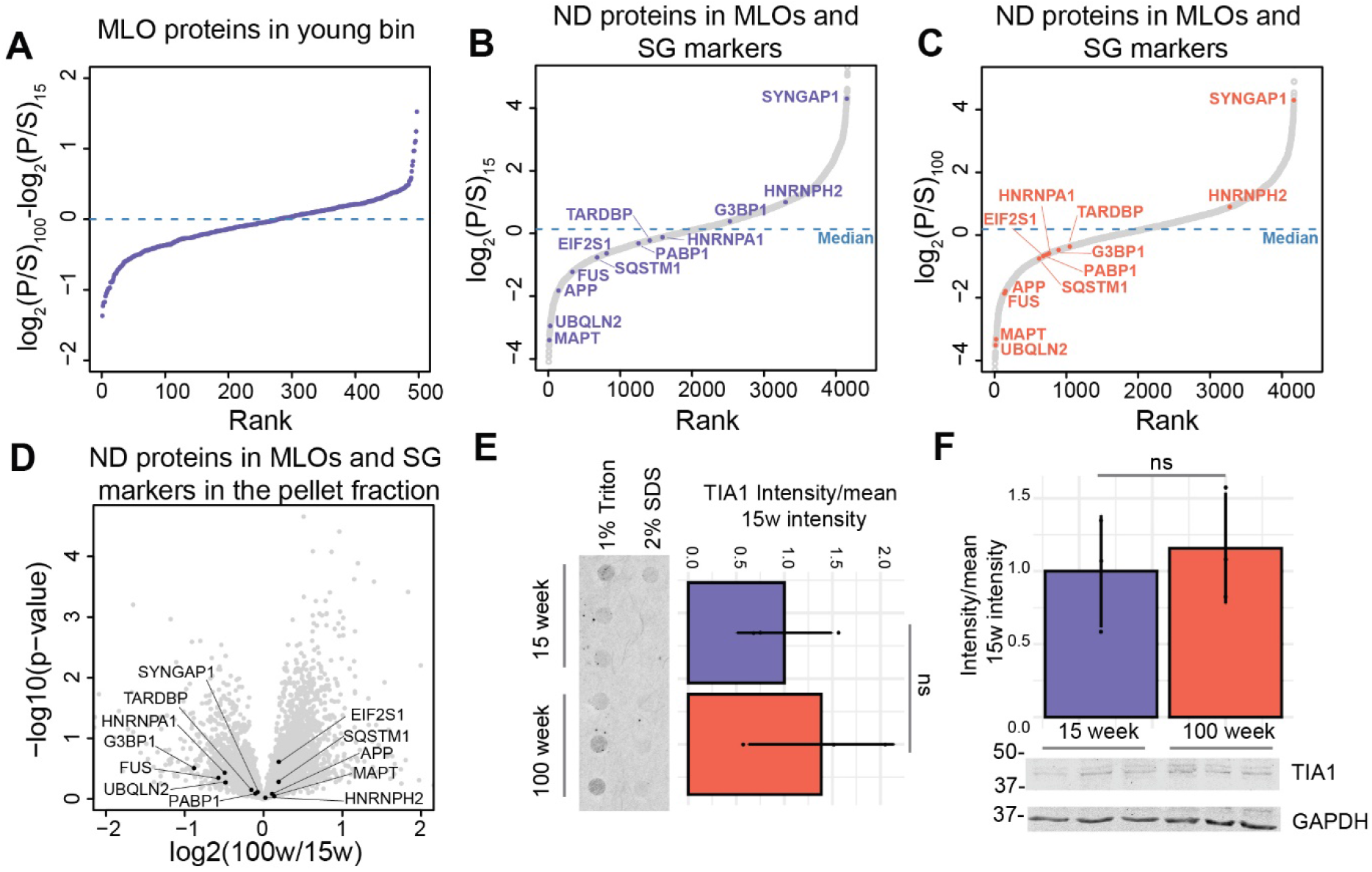
MLO proteins are found in the pellet fraction of young and old mice. A) Ranked plot of the difference in pellet/supernatant ratio between old and young mice for MLO proteins enriched in the pellet fraction in the combined young datasets. B-C) Ranked plot of the average P/S ratio of all quantified proteins in young (B) and old (C) mice. Selected disease-associated proteins that localize in condensates and/or stress granules are indicated. D) Selected proteins are highlighted in the volcano plot that compares composition of detergent insoluble fractions obtained from the cortex of 15 and 100-week-old (same data shown in Figure 1B). E) The FTA signals are normalized to the mean of the corresponding young data points. F) Western blot signal from TIA1 is normalized the GAPDH loading control and then normalized to the mean of the young data points.

**Figure S8.**
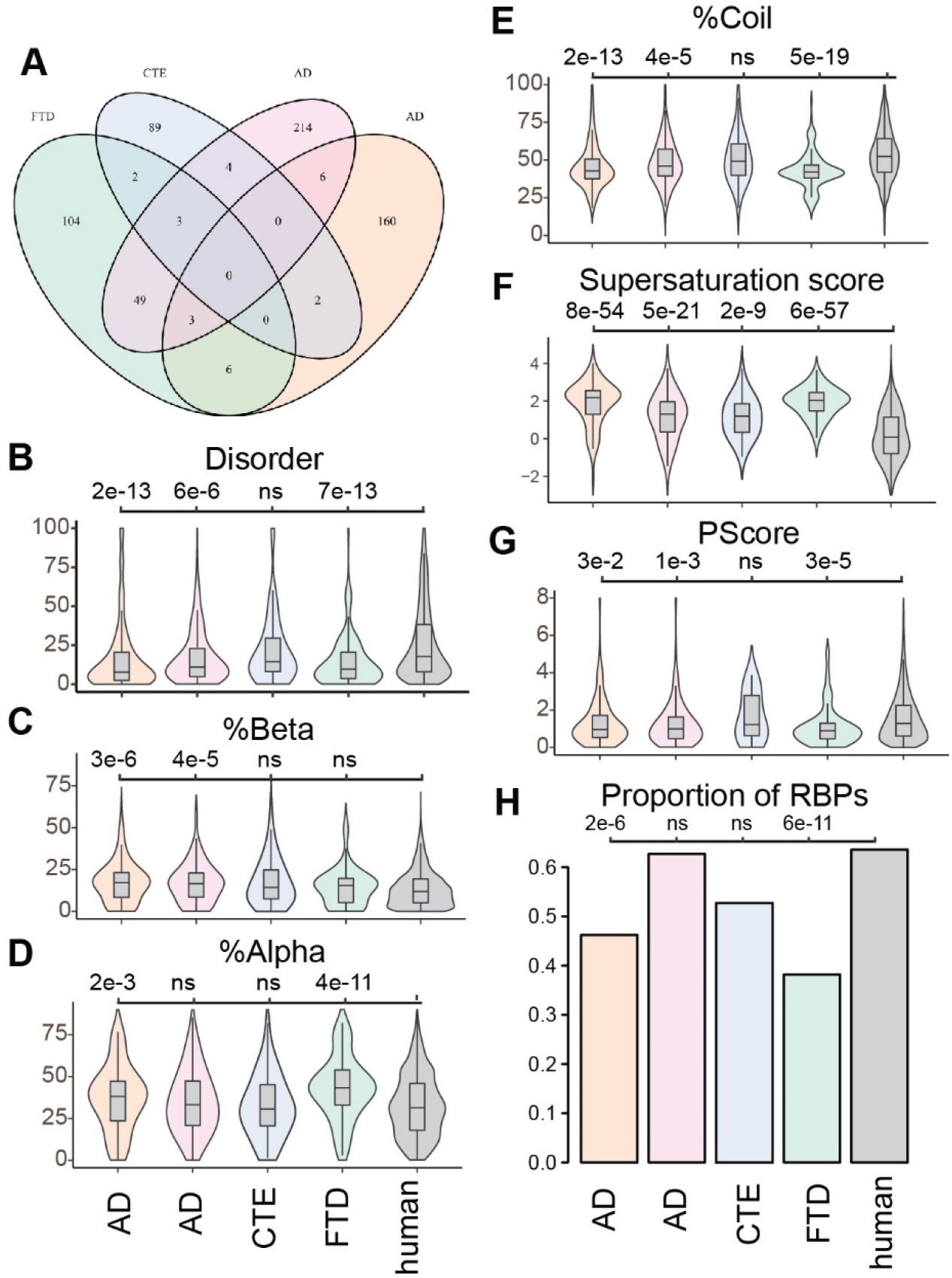
Human solubility datasets share similar features to aged mice. A) Venn diagram representing the overlap between the selected human MS datasets; B-G) Violin plots comparing the distribution of proteins with the indicated features including: B) percentage of protein sequences predicted to be intrinsically disordered; C) percentage of sequence in beta sheet; D) percentage of sequence alpha helix; E) percentage of sequence coils; F) Each protein’s supersaturation score. G) P-Scores indicating the propensity of each protein to participate in pi-pi interactions. P-values for all violin plots were calculated using a Hochberg adjusted Wilcoxon test. H) The proportion of proteins in each dataset predicted to RNA-binding proteins (RBPs). Fisher test followed by Hochberg correction.

